# Subliminal beauty engages the brain’s valuation circuits

**DOI:** 10.1101/2025.11.17.688735

**Authors:** Patrícia Fernandes, Joseph W. Kable, Jorge Almeida, Anita Tusche, Christian C. Ruff, Fredrik Bergström

## Abstract

Neuroeconomic models propose that the anterior ventral striatum (aVS) and ventromedial prefrontal cortex (vmPFC) are key regions for computing subjective value (SV) signals that guide choice. However, the role of conscious awareness in this process remains debated. Here we examined whether SV can be automatically computed in these regions without conscious awareness. In an fMRI experiment, participants viewed faces that varied in attractiveness under three conditions: (i) suppressed from awareness via continuous flash suppression (CFS), (ii) clearly visible without suppression, or (iii) absent (background only) with CFS. Participants reported trial-wise facial identity (objective) and visibility (subjective) measures of facial awareness. In a post-fMRI session, they rated the attractiveness of each face as a measure of SV. When faces were seen, task performance (d’) exceeded chance and responses were faster than in absent trials. When faces were unseen, performance was at chance but responses were slower than absent trials. Neurally, seen and unseen faces elicited greater neural signal than absent trials and showed similar neural patterns in the fusiform face area (FFA). Critically, neural signal in vmPFC correlated with SV for both seen and unseen faces, with similar neural patterns. In aVS, SV-related signal was only observed for unseen faces. Furthermore, mean face-related signal in FFA correlated with SV-related signals in aVS and vmPFC for unseen faces. These findings demonstrate that SV can be automatically computed in aVS and vmPFC without conscious awareness, suggesting a neural pathway by which subliminal information can influence value-based choice.

## Introduction

We may think we are aware of all the information influencing our decisions. However, this intuition is likely misleading since we are only consciously aware of a fraction of all neural processes at any given time. Many neural processes may operate automatically below our conscious awareness as we go about our lives. For instance, our brain seems to automatically extract useful information from our consciously perceived environment, such as affordances and other object properties when seeing manipulable objects (Almeida et al., 2023; Bergström et al., 2021; Valério et al., 2025); and trustworthiness, dominance features, and attractiveness when seeing faces (Engell et al., 2007; Kim et al., 2007; Lebreton et al., 2009; Miao et al., 2022). To what extent decision processes operate outside of our conscious awareness and influence our choices is not well understood and heavily debated (Bechara et al., 1997; Libet et al., 1983; Maia & Mcclelland, 2004; Maoz et al., 2019; Newell & Shanks, 2014; Schurger et al., 2012; Soon et al., 2008). This is especially true for value-based decisions, which involve evaluating and comparing options based on their subjective value (SV), an individualized estimate of their desirability or worth. This process enables organisms, from nematodes to humans, to adapt their actions to maximize perceived benefit and underlies human behaviors such as consumer purchases, mate selection, and moral judgments. We tested whether SV, operationalized as facial attractiveness, can be automatically computed in the brain’s valuation circuits when faces are suppressed from conscious awareness with continuous flash suppression (CFS).

Explicit evaluation of consciously presented objects (e.g., faces, food, trinkets, etc.) is consistently associated with a positive relationship between their SV and neural signals in aVS and vmPFC (Chatterjee et al., 2009; Chib et al., 2009; Lebreton et al., 2009; Pegors et al., 2015). Because these core valuation circuits represent the SV of choice options across stimuli and tasks, their activity is thought to reflect an abstract “common currency” that enables the brain to compare diverse options, such as food, money, or social approval, by translating them into a shared value scale that guides choice (Bartra et al., 2013; Chib et al., 2009; Dang et al., 2024; Kable & Glimcher, 2007, 2009; Kobayashi & Hsu, 2019; D. J. Levy & Glimcher, 2012; I. Levy et al., 2010; McNamee et al., 2013; Pegors et al., 2015; Shuster & Levy, 2018). Consistently, individual variation in vmPFC anatomy (e.g., grey matter volume) or vmPFC lesions have been associated with individual differences or disruptions of choice preferences (Bergström, Lerman, et al., 2024; Bergström, Schu, et al., 2024; Camille et al., 2011; Fellows & Farah, 2007; Godefroy et al., 2024; Pehlivanova et al., 2018; Yu et al., 2022).

Interestingly, neuroimaging research has demonstrated that SV computations in the brain can arise automatically, even without explicit evaluation instructions, by presenting consciously perceived objects during passive viewing or while performing unrelated distractor tasks. These studies found that neural signals or patterns in aVS and vmPFC correlated with SV and could predict real or hypothetical choice (Kim et al., 2007; Lebreton et al., 2009; Levy et al., 2011; Smith et al., 2014; Tusche et al., 2010), while other studies showed that SV signals in these regions are strongly enhanced when SV is explicitly used to guide choice rather than when computed automatically during an unrelated distracting task (Grueschow et al., 2015). A key limitation is that when stimuli are consciously presented, it is difficult to rule out the influence of spontaneous conscious evaluations, as these cannot be directly measured without potentially inducing them. This limitation can be addressed by presenting stimuli subliminally, thereby eliminating the possibility of conscious evaluation.

Using subliminal stimuli to investigate neural processes without conscious influence introduces other methodological challenges, and studies examining SV processing under such conditions are scarce with mixed results. For example, subliminal cues linked to monetary rewards or punishments modulated aVS signal and influenced choice, suggesting subliminal valuation and learning (Pessiglione et al., 2008). Similarly, subliminal erotic compared to neutral images engaged the aVS, consistent with automatic processing of biologically relevant value (Oei et al., 2012). In contrast, other studies using subliminal images of coins (Bijleveld et al., 2014; Pessiglione et al., 2007), erotic images (Gillath & Canterberry, 2012), or faces with varying attractiveness (Ito et al., 2015), did not find subliminal SV effects in aVS or vmPFC. Instead, when subliminal effects were observed, they appeared outside core valuation circuits and likely reflected other processes than SV. Because reported subliminal SV effects were confined to subcortical areas, it was suggested that conscious awareness is necessary for SV processing in the prefrontal cortex (Bijleveld et al., 2012).

However, none of those studies collected trial-wise awareness measures during their main experiments. Instead, awareness was inferred from group-level discrimination performance (at chance) before or after the task, which risks including trials with conscious awareness. This may explain why Pessiglione et al.’s (2008) findings were not replicated in a behavioral study that applied trial-wise awareness measures (Skora et al., 2023). Moreover, those studies used backward masking with very brief stimuli durations that may not yield as strong subliminal effects as other methods such as CFS or attentional blink, which may explain the mixed results.

We recently challenged the view that prefrontal SV processing requires conscious awareness, by showing that subliminal probabilities could be integrated with consciously perceived rewards into SV in both aVS and vmPFC during risky choice, using trial-wise awareness measures in an attentional blink paradigm (Fernandes et al., 2025). However, because the subliminal information was directly relevant to the task, its processing may have depended on top-down task goals and demands rather than purely automatic bottom-up mechanisms that are independent of task goals or demands.

Here, we investigated whether SV, operationalized as a continuous measure of facial attractiveness, can be automatically computed in aVS and vmPFC without conscious awareness. We used CFS to present faces subliminally or consciously, used trial-wise face-identification and subjective awareness reports, and tested for a positive relationship between neural signal in the core valuation circuits and post-fMRI SV ratings. We found that subliminal faces elicited SV signals in both aVS and vmPFC, indicating that SV computations can occur automatically and operate independently of conscious awareness.

## Materials and Methods

### Participants

We recruited 36 healthy volunteers from the University of Coimbra campus area. All participants had normal or corrected-to-normal vision, gave written informed consent, received extra course credits (if applicable), and a 20-euro reward voucher for participation. The study was approved by the Ethics Committee of the Faculty of Psychology and Educational Sciences of the University of Coimbra, Portugal.

Four participants were excluded prior to the fMRI session due to more than 20% suppression breaks during practice. Following the fMRI session, one participant was excluded for unusually high task performance (d’) when faces were reported unseen (d’ > 3 SD) to avoid the possibility of conscious influences; one for irretrievable MRI data corruption; and one for insufficient unseen trials caused by frequent suppression breakthroughs (17 trials in total for all runs). In addition, one run was excluded from a participant who had only one unseen trial in that run. We therefore used 29 participants (24 females; 17 – 28 age range, M = 20, SD = 2.7 years; 3 with same-sex attraction) for our analyses.

### Stimuli and Procedure

The study consisted of a 30-45 min instruction and practice session (Mdn = 28 days) prior to a 90 min fMRI session that was immediately followed by post-fMRI attractiveness ratings. The fMRI task was programmed in OpenSesame, stimuli (400 x 400 pixels, roughly subtending 7.27 degrees visual angle) was presented on a BOLDscreen 32UHD (60 Hz, 1024 x 768 resolution) (Cambridge Research Systems Ltd., UK), and responses were collected with a Cedrus Lumina button box. Stimuli were viewed through red–green anaglyph glasses to enable continuous flash suppression (CFS) during the face-identification task.

The fMRI task had 261 trials in total across three stimulus presentation conditions (Figure 1A): (i) a clearly visible face without CFS, presented in gray-scale to both eyes, superimposed on green background and moving red Mondrians (n = 72; Face + No CFS); (ii) a face rendered invisible by CFS, a green-scaled face presented to the non-dominant eye, and moving red Mondrians to the dominant eye (n = 144; Face + CFS); and (iii) absent face, a uniform green background presented to the non-dominant eye, and moving red Mondrians to the dominant eye (n = 45; No Face + CFS). When faces were presented to both eyes, they were always consciously perceived. When faces (or empty backgrounds) and Mondrians were presented to different eyes, only the Mondrians were consciously perceived, whereas the faces or backgrounds were suppressed but subliminally processed in the brain, but occasionally broke through suppression. Following prior work (Bergström & Eriksson, 2018; Fontan et al., 2021; Pedale et al., 2023), trials were categorized as seen (visible and reported seen), unseen (suppressed and reported unseen), or absent (absent and reported unseen) based on presentation condition and PAS reports.

**Figure 1.**
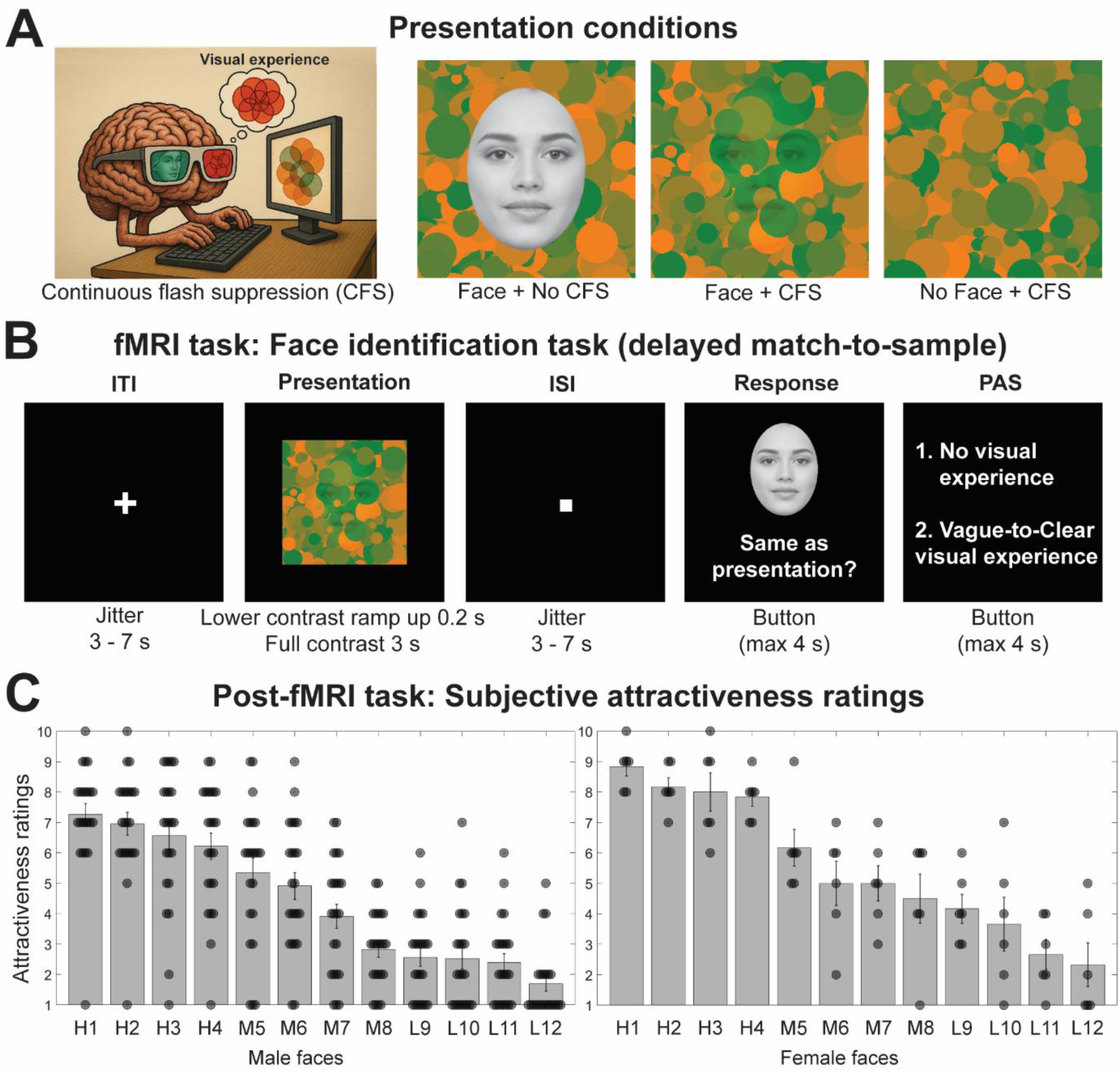
Experimental procedure. (**A**) The experiment had three presentation conditions: a face suppressed by continuous flash suppression (CFS; Face + CFS), a uniform background (absent a face) suppressed by CFS (No Face + CFS), and a clearly visible face without suppression (Face + No CFS). The image of the brain with anaglyph glasses is a schematic illustration of the CFS setup and participants visual experience, and was made with ChatGPT4 (**B**) fMRI delayed match-to-sample task procedure: Jittered inter-trial-interval (ITI), face presentation, jittered inter-stimulus-interval (ISI), sample-to-match response (counterbalanced buttons for responses), perceptual awareness scale (PAS). (**C**) Post-fMRI task: Mean subjective attractiveness ratings for each face used in the experiment. Participants were only presented with images of faces from the sex they were most attracted to. The faces are numbered from 1 to 12 from highest to lowest mean ratings, and prefixed with a letter that indicates whether that face was belonged to the preselected high, medium, or low attractiveness category. The face in the illustrations (A) and (B) is AI generated and used with permission from Generated Photos (https://generated.photos).

The original face images were cropped in an oval shape to exclude non-facial features, Gaussian blur (1 pixel radius) was applied, the opacity of the face was lowered to blend in with the background, additional blur was applied to the edge around the face, the SHINE Toolbox (https://github.com/RodDalBen/SHINE_color; Dal Ben, 2023) was used to match luminance and spatial frequency while maintaining the structure of the faces, and custom Matlab code was used to make the modified face images and empty backgrounds green and to create red Mondrian images to be used for CFS (http://martin-hebart.de/webpages/code/stimuli.html; Stein et al., 2011).

Each trial (Figure 1B) consisted of: a jittered fixation cross (3 - 7 s), stimulus presentation (3 s; suppressed faces ramped in over 200 ms to reduce suppression breaks), jittered fixation dot (3 - 7 s), a match-to-sample response (match or non-match; max 4 s), perceptual awareness scale (PAS) report (1 = No visual experience of a face, 2 = vague-to-clear visual experience of a face; max 4 s; Fernandes et al., 2025; Sandberg et al., 2010). Participants were instructed to memorize the face if they saw it, and after the delay, indicate whether the sample face matched the previously presented face. They were told that a face was always presented, but mostly suppressed, and that when they did not see it, they should guess based on their gut-feeling or intuition rather than rely on systematic strategies (e.g., always pressing “non-match”). After each trial, they provided their PAS rating for the face. Importantly, the task contained no attractiveness judgments, and participants were unaware that they would later provide post-fMRI attractiveness ratings.

Immediately after scanning, participants rated the attractiveness of all 12 unique faces from the fMRI session on a 1–10 scale via Google Forms. Ratings were unrestricted in time and participants were told they could reuse ratings across faces. These ratings served as our measure of SV.

The twelve unique faces used in the experiment were pre-selected to represent three categories of attractiveness (low, medium, high), to ensure a wide distribution of subjective ratings across participants. All faces appeared in both visible and suppressed conditions, with more repetitions in the suppressed condition. Faces in the low and medium categories were taken from the Chicago Face Database (Ma et al., 2015), corresponding to the lowest and mid-range average ratings. Because the database did not include faces at the top of the attractiveness scale, we supplemented our stimulus set with AI-generated model faces (https://generated.photos) for the high-attractiveness category.

This approach ensured coverage across the full attractiveness spectrum and maximized variance for the parametric SV analyses. Participants viewed only faces of the sex they reported being most attracted to, as indicated in a pre-screening form that also included demographic, eligibility, and MRI safety questions. The reported attractiveness ratings confirmed a wide spread across the full scale, with substantial individual variability in both male and female stimulus sets (Figure 1C; individual ratings in SI-Figure 1).

Before being scheduled for the fMRI session, participants completed a shorter practice version of the task to ensure that they understood the instructions and that continuous flash suppression (CFS) worked effectively on them. Task instructions were first provided in writing and then explained in detail by a researcher before the practice began. To secure a sufficient number of unseen trials for later analyses, only participants who saw fewer than 20% of faces during practice were invited to proceed to the fMRI session.

### MRI acquisition

MRI data were acquired on a 3T MAGNETOM Vida whole-body scanner (Siemens Healthineers, Erlangen, Germany) at Hospital da Luz de Coimbra using a 32-channel head coil. Each participant completed one scanning session consisting of three functional runs (746 volumes per run; 2238 volumes and 75 min in total) with a high-resolution anatomical scan acquired between runs.

Anatomical MRI images were acquired using a T1-weighted magnetization-prepared rapid gradient echo (MPRAGE) sequence [repetition time (TR) = 2300 ms, echo time (TE) = 2.32 ms, voxel size = 0.9 x 0.9 x 0.9 mm^3^, flip angle = 8 degrees, field of view (FoV) = 240 x 240, matrix size = 256 x 256, bandwidth (BW) = 200 Hz/px, GRAPPA acceleration factor 2].

Functional MRI (fMRI) data were acquired using a T2*-weighted gradient echo planar imaging (EPI) sequence (TR = 2000 ms, TE = 36 ms, voxel size = 3 x 3 x 3 mm^3^, slice thickness = 3 mm, FoV = 210 x 210, matrix size = 70 x 70, flip angle = 75 degrees, BW = 1786 Hz/px). Each volume contained 50 contiguous transverse slices acquired in interleaved order, oriented parallel to the AC–PC line and tilted 30° to improve signal coverage in the vmPFC.

### Preprocessing of MRI data

MRI data were preprocessed with fMRIPrep 24.0.1 default settings (see Supplemental information for a full detailed description). For anatomical images, the T1-weighted scan was corrected for intensity non-uniformity, skull-stripped, segmented into gray matter, white matter, and CSF, and spatially normalized to MNI space via non-linear registration. For functional images, preprocessing included generation of a reference volume (custom fMRIPrep method), head-motion correction (six rigid-body parameters), and slice-time correction to middle slice. The BOLD reference was co-registered to the T1 reference image. Preprocessed BOLD series were then normalized to MNI space. 24 motion parameters, several confounding time series, and physiological noise regressors were automatically calculated by fMRIprep defaults. Finally, an 8 mm FWHM Gaussian kernel was applied with SPM12 before statistical analysis.

### Statistical analysis

As in previous work (Bergström & Eriksson, 2018; Fontan et al., 2021; Pedale et al., 2023), we defined condition categories in the following way: “Seen” was defined as a visibly presented face without suppression and participants reporting a visual experience of the face (PAS = 2). “Unseen” was defined as a face presented with suppression and participants reporting no visual experience of the face (PAS = 1). “Absent” was defined as the absence of a face presented with suppression and participants reporting no visual experience of a face (PAS = 1). These three seen, unseen, and absent categories and naming convention are used for all the subsequent analysis. Trials not included in these categories were grouped into an “other” condition for fMRI modeling but otherwise not of interest.

#### Behavioral analysis

We looked at task performance and reaction time. Task performance (d’) was defined as z(hits) minus z(FAs), where “hits” was when the to be remembered face matched the sample face and the response was “match”, while “false alarms” was when the to be remembered face did not match the sample and response was “match.” Prior to reaction time analysis, we removed outlier trials with a reaction time < 250 ms (e.g., accidentally pressing the button before seeing the query) or > three standard deviations from the group mean (e.g., spacing out), and log10-transformed the reaction times for a normal distribution (Ratcliff, 1993).

#### fMRI analysis

##### General Linear Model (GLM)

Within-participant modeling was conducted in SPM12 using a GLM with restricted maximum likelihood estimation. The model included 14 regressors of interest: unseen face presentation, unseen linear parametric modulation of SV, unseen ISI, unseen response; seen face presentation, seen linear parametric modulation, seen ISI, seen response; absent face presentation, absent ISI, absent response; other face presentation, other ISI, other response. The “other” category contained trials that did not fit the seen, unseen, and absent categories (e.g., faces breaking suppression).

Parametric SV regressors for seen and unseen faces were modeled as linear modulations, using run-wise mean-centered and z-scored subjective attractiveness ratings from the post-fMRI session. Regressors of interest were modeled from stimulus onset to offset and convolved with the canonical hemodynamic response function. A high-pass filter with a cut-off of 128 s and a global AR(1) autocorrelation model were applied.

The GLM also contained 40 regressors of no interest: 24 head motion parameters, 4 cerebrospinal fluid (CSF) CompCor components, 11 white matter CompCor components, and one framewise displacement parameter. The number of CompCor components was equalized across participants and runs by selecting the maximum number of components available for all participant’s runs (Fernandes et al., 2025).

For multivariate analyses, a separate GLM was computed to estimate beta maps. The model included 46 regressors of interest per run: one regressor for each of the unique unseen faces presented (12 in total), one regressor for each unique seen face presented (12 in total), one regressor for each absent presentation (15 in total), ISI and response regressors separated by seen, unseen, and absent conditions (6 in total), and a “other” regressor. No parametric modulation regressors. The same 40 regressors of no interest to model motion and physiological noise as before. This procedure therefore yielded up to 12 beta maps for unseen and 12 beta maps for seen face presentations per run (36 seen and 36 unseen total per participant), as well as 15 beta maps for absent trials (45 total per participant).

##### Regions of interest

We focused on three a priori regions of interest (ROIs) based on previous literature. The anterior ventral striatum (aVS) and ventromedial prefrontal cortex (vmPFC) were selected because they form part of the brain’s core valuation circuit, consistently associated with subjective value (SV) but not salience (for meta-analysis, see Bartra et al., 2013) and reliably engaged by facial attractiveness (for meta-analysis, see Chuan-Peng et al., 2020). The fusiform face area (FFA) was included as a high-level visual face processing region (Grill-Spector et al., 2017; Kanwisher et al., 1997).

The aVS was defined using the AAL3 anatomical atlas to ensure precise coverage of the entire region while avoiding neighboring structures, and was used for both univariate and multivariate analyses. The vmPFC was defined by placing a spherical ROI centered on the coordinates [-1, 46, −7] from Bartra et al.’s (2013) SV meta-analysis. For univariate analyses, we used a 4 mm radius sphere, while for multivariate analyses we used a 12 mm radius sphere. The FFA was defined using peak coordinates [40, −50, −20] from a NeuroSynth meta-analysis with the search term “faces,” with spheres of 4 mm radius for univariate and 12 mm for multivariate analyses. We removed parts of the larger FFA spheres that crossed over to the cerebellum. We used smaller spheres in univariate analyses to target peak effects as precisely as possible, and larger spheres for the multivariate analyses to capture distributed voxel patterns, with the cross-validation procedure automatically selecting the subset of voxels most predictive within each sphere, and thus using the two approaches in a complimentary way.

##### Univariate analysis

For ROI analyses, the mean contrast estimate across all voxels in the ROI were extracted for each participant and entered into group-level t-tests. One-tailed tests were used whenever we had strong a priori hypotheses derived from previous literature (e.g., neural responses in valuation circuits should scale positively with SV), or when only one direction of the effect was meaningful (e.g., neural signal or decoding being greater than chance).

To test whether stronger face-related signal in the FFA was related to stronger SV-related signal in valuation regions during the face presentation, we correlated contrast estimates for seen faces > absent in FFA with the parametric SV estimates for seen faces in aVS and vmPFC, as well as, unseen faces > absent in FFA with the parametric SV estimates for unseen faces in aVS and vmPFC.

In addition to ROI analyses, we performed whole-brain contrasts to test for effects outside our a priori regions. These analyses were restricted to a grey matter mask and evaluated using non-parametric permutation testing (10,000 permutations) with Threshold-Free Cluster Enhancement (TFCE; http://www.neuro.uni-jena.de/tfce/; Smith & Nichols, 2009) to adjust for cluster size and voxel-wise FDR correction for multiple comparisons.

##### Multivariate pattern analysis

To assess out-of-sample predictions (or “decoding”), we used a Thresholded Partial Least Squares regression (T-PLS; https://github.com/sangillee/TPLSm; Lee et al., 2022) implemented in MATLAB R2022a. T-PLS was selected because it can flexibly predict both continuous and binary variables, handles high voxel collinearity, performs voxel selection within a nested cross-validation framework, is computationally efficient, and provides interpretable coefficients. We performed two main sets of analyses: (i) binary decoding (seen face vs. absent; unseen face vs. absent; coded as 1 for face and 0 for absent) to test for face-related information in the FFA, and (ii) SV decoding as a continuous variable (for seen and unseen faces) to test for SV signal in aVS and vmPFC.

We applied a leave-one-participant-out cross-validation (LOPO-CV) procedure with a nested LOPO-CV inside each training set to select optimal tuning parameters, number of PLS components (1 – 15) and proportion of voxels retained (0 – 100% in 25% steps), based on highest prediction accuracy. After parameter optimization, the model was fit to the training data and tested on the left-out participant. For binary face decoding, the training data consisted of approximately 2,268 beta maps (28 participants × 3 runs × 27 beta maps) and the test set of 81 beta maps (1 participant × 3 runs × 27 beta maps) per fold. Binary prediction accuracy was quantified using Area Under the Curve (AUC). For continuous SV prediction, the training set comprised approximately 1,008 beta maps (28 participants × 3 runs × 12 beta maps) and the test set 36 beta maps (1 participant × 3 runs × 12 beta maps) per fold, with prediction accuracy defined as Pearson’s correlation between predicted and observed SV values.

Statistical significance of decoding performance was evaluated using permutation testing. Labels were randomly shuffled within each participant and run, and the analysis was repeated 10,000 times to generate a null distribution. One-tailed p-values were computed by comparing the observed median prediction accuracy to the null distribution. The combined use of AUC metrics and permutation testing ensured that any slight category imbalance (12 face vs. 15 absent trials) did not bias the face decoding results.

To generate a null distribution for pair-wise comparisons between seen face vs. absent and unseen face vs. absent decoding accuracies, we applied a sign-flip procedure. On each of the 10,000 permutations, the sign of each participant’s accuracy difference was randomly flipped (equivalent to randomly swapping condition labels within participants), and the median of the permuted differences was calculated. This procedure preserved the within-participant dependency structure and yielded a null distribution that reflected the absence of a group difference. The two-sided p-value was calculated as the proportion of permuted statistics with absolute values greater than or equal to the observed statistic.

We also performed cross-decoding to test generalization across conscious awareness (seen and unseen conditions). Using the same LOPO-CV framework, we trained the model on seen face beta maps and tested it on unseen faces in the left-out participant, and vice versa, then averaged prediction accuracy across directions. Importantly, none of the left-out-participant’s data was used in the training fold to avoid any statistical dependency between training and testing data. For null distributions, cross-decoding accuracies were likewise averaged across directions before computing permutation-based p-values.s

Finally, to generate interpretable T-PLS model coefficients, we built a separate predictor model following the LOPO-CV framework but without nested parameter optimization. This approach allowed us to fit the model to all participants’ data and extract voxel-level coefficients highlighting predictive patterns for both binary and continuous out-of-sample decoding.

## Results

### Behavioral results

#### Perceptual awareness scale ratings

To confirm that the CFS paradigm worked as intended, we examined the perceptual awareness scale (PAS) responses for the faces in visible, suppressed, and absent conditions (Figure 1 B). When faces were visibly presented without suppression, they were reported as seen (PAS = 2) on most trials (Mdn _proportion_ = 95.8%, SE _proportion_ = 2.3%, Mdn _trials_ = 69, SE _trials_ = 0.02) with only a few mistakes. When faces were suppressed by CFS, they were reported as unseen (PAS = 1) on most trials (Mdn _proportion_ = 81.3%, SE _proportion_ = 4.2%, Mdn _trials_ = 117, SE _trials_ = 0.04) but sometimes breaking suppression. When faces were absent and an empty background was suppressed with CFS, most trials were reported as unseen (Mdn = 97.8%, SE = 2.6%, Mdn = 44, SE = 0.03), with few false alarms. Overall, the CFS paradigm worked well, awareness reports aligned with expectations and we had enough trials for analysis.

#### Task performance and reaction times

On trials where the face was “seen” (clearly visible and reported seen), task performance (d’) was much greater than chance (M = 3.37, SE = 0.15, t(28) = 23.08, p < .001, one-tailed; SI-Figure 2A). However, when faces were “unseen” (suppressed and reported unseen), task performance (d’) was not greater than chance (M = 0.06, SE = 0.06, t(28) = 0.93, p = .18, one-tailed). As expected, task performance for seen faces was much greater than for unseen faces (t(28) = 20.74, p < .001, two-tailed). It was not possible to compute task performance for absent trials since there was no face. There was no direct influence of the unseen faces on task performance, and faces can therefore be considered subliminal by both objective and subjective measures of awareness.

When testing for indirect influences, we examined whether trials with seen and unseen faces differed from trials absent faces, and whether there was a difference between unseen trials when responses were correct (hits & correct rejections) and incorrect (misses & false alarms). Because the face-identification task was easy when the faces were seen, seen incorrect trials were only present in 17 of 29 participants, we therefore did not include them in the ANOVA. The 1 x 4 repeated measures ANOVA revealed a difference (F(3,28) = 14.76, p < .001) in reaction time across: seen correct (M_log(RT)_ = 3.06, SE = 0.01), absent (M_log(RT)_ = 3.11, SE = 0.01), unseen correct (M_log(RT)_ = 3.12, SE = 0.02), and unseen incorrect (M_log(RT)_ = 3.13, SE = 0.01).

Post hoc t-tests (SI-Figure 2B) revealed that reaction times were faster for seen correct compared to absent (t(28) = −3.26, p_FDR_ = .006, p_unc._ = .003, two-tailed); unseen correct (t(28) = −4.09, p_FDR_ = .001, p_unc._ < .001, two-tailed); and unseen incorrect (t(28) = −4.65, p_FDR_ < .001, p_unc._ < .001, two-tailed). Interestingly, reaction time for unseen correct (t(28) = −2.58, p_FDR_ = .019, p_unc._ = .015, two-tailed) and unseen incorrect (t(28) = −3.11, p_FDR_ = .006, p_unc._ = .004, two-tailed) responses were both slower than absent trials. However, there was no difference between unseen correct and incorrect trials (t(28) = −0.41, p_FDR_ = .685, p_unc._ = .685, two-tailed).

Taken together, these results show that even though unseen and absent faces had the same lack of perceptual awareness and unseen task performance was at chance, the unseen faces slowed reaction time compared to absent trials, suggesting a subliminal memory trace of the unseen faces.

### fMRI results

In the sections below, we first present the univariate results, which test whether the mean neural signal differs between conditions, followed by the multivariate decoding results, which assess whether neural patterns differ between conditions, or in the case of cross-decoding, whether patterns are similar across conditions. For brevity, these results are presented together according to section topic because they are both testing for the presence of face or SV processing. As a reminder, we used the same condition definitions as in the behavioral analyses: “seen” (clearly visible face and reported seen), “unseen” (suppressed face and reported unseen), and “absent” (no face and reported unseen).

#### Face perception in FFA

To confirm that both seen and unseen faces were processed in higher-level visual areas, we conducted univariate and multivariate ROI analyses in the fusiform face area (FFA; Figure 2).

**Figure 2.**
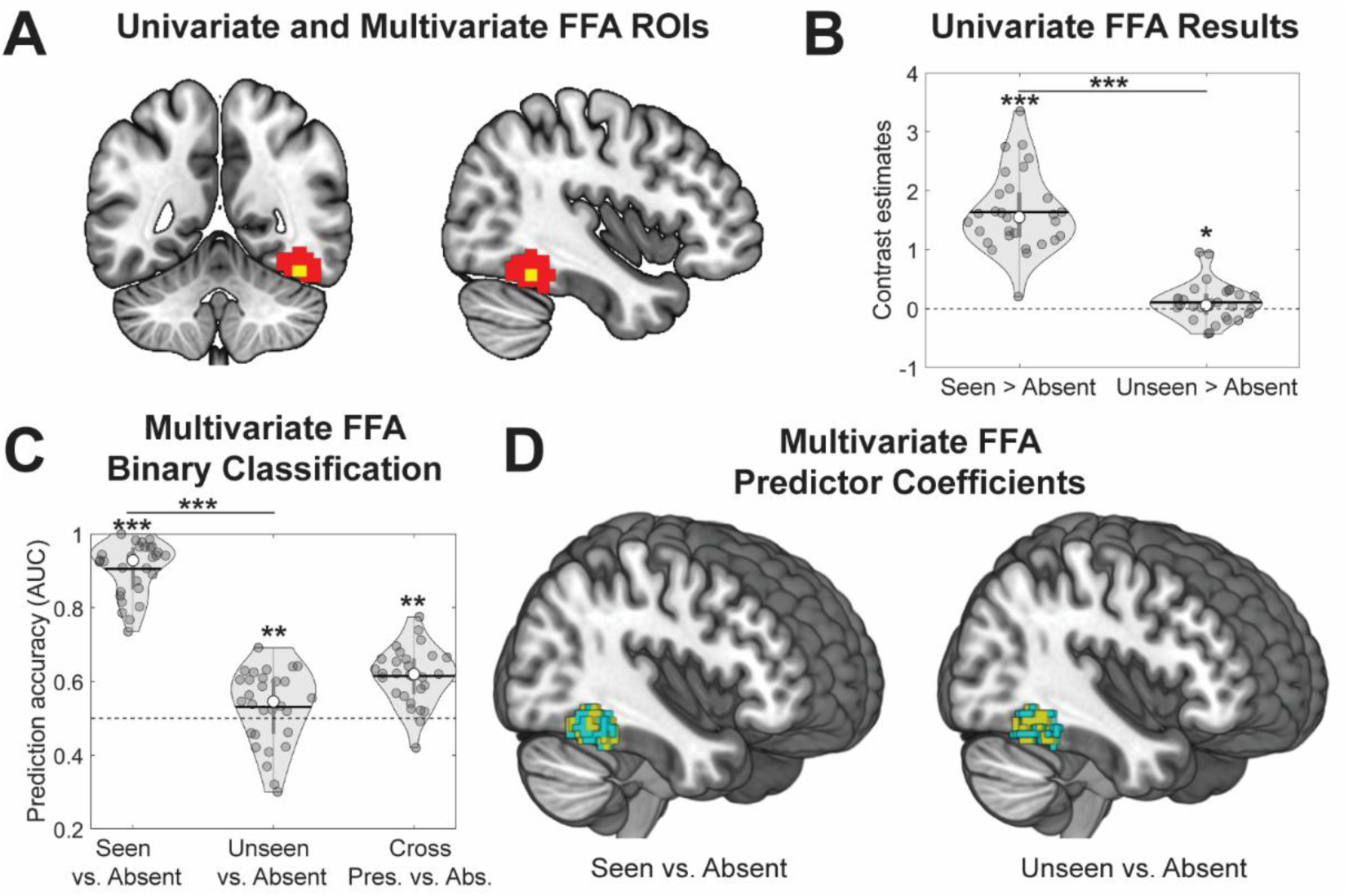
Face perception. (**A**) Regions of interest (ROI) for univariate (yellow) and multivariate (red) analyses for the fusiform face area (FFA). (**B**) Univariate FFA results for contrasts seen > absent and unseen > absent. (**C**) Multivariate classification results for seen vs. absent, unseen vs. absent, and cross-decoding presence vs. absence across seen and unseen faces. (**D**) Predictor coefficients for the subset of voxels that contributed to significant multivariate face decoding results. Warm color indicates a positive, and cold color a negative, relationship with neural signal and the presence of a face. P-values were FDR-adjusted separately within each test set: 5 tests against chance and 2 paired tests. * p < .05, ** p ≤ .01, *** p ≤ .001.

##### Seen faces

As expected, seen > absent faces showed significant mean neural signal change (M = 1.62, SE = 0.12, t(28) = 13.20, p_FDR_ < .001, p_unc._ < .001, one-tailed), and seen vs. absent out-of-sample classification showed significant prediction accuracy (Mdn_AUC_ = 92.8, p_FDR_ < .001, p_unc._ < .001, one-tailed permutation test), with a mix of positive and negative predictor coefficients in the FFA.

##### Unseen faces

Similarly, unseen > absent faces showed significant neural signal change (M = 0.10, SE = 0.06, t(28) = 1.76, p_FDR_ = .045, p_unc._ = .045, one-tailed), and unseen vs. absent classification showed a significant prediction accuracy (Mdn_AUC_ = 54.6, p_FDR_ = .006, p_unc._ = .005, one-tailed permutation test),s with a mix of positive and negative predictor coefficients in the FFA.

##### Cross-decoding

Presence vs. absence could be cross-decoded across seen and unseen faces in the FFA (Mdn_AUC_ = 62.0, p_FDR_ = .002, p_unc._ < .001, one-tailed permutation test). Taken together, these findings suggest that both seen and unseen faces were processed in the FFA with a shared neural code.

##### Seen and unseen comparisons

The neural difference for seen > absent faces were significantly greater than unseen > absent faces (t(28) = 13.79, p_FDR_ < .001, p_unc._ < .001, two-tailed), and the decoding accuracy for seen vs. absent faces was significantly greater than unseen vs. absent faces (p_FDR_ < .001, p_unc._ < .001, two-tailed permutation test). These results show that seen faces produced a much stronger neural response in the FFA than unseen faces.

Together, these results demonstrate that unseen faces were processed with a similar but weaker neural representation than seen faces in FFA.

#### Subjective value in aVS and vmPFC

To test whether subjective value (SV; operationalized as the idiosyncratic continuous measure of overall facial attractiveness) can be automatically computed in the aVS and vmPFC independently of conscious awareness, we tested for a positive relationship between SV ratings and neural signal for both seen and unseen faces with univariate parametric modulations and multivariate regression predictions (Figure 3).

**Figure 3.**
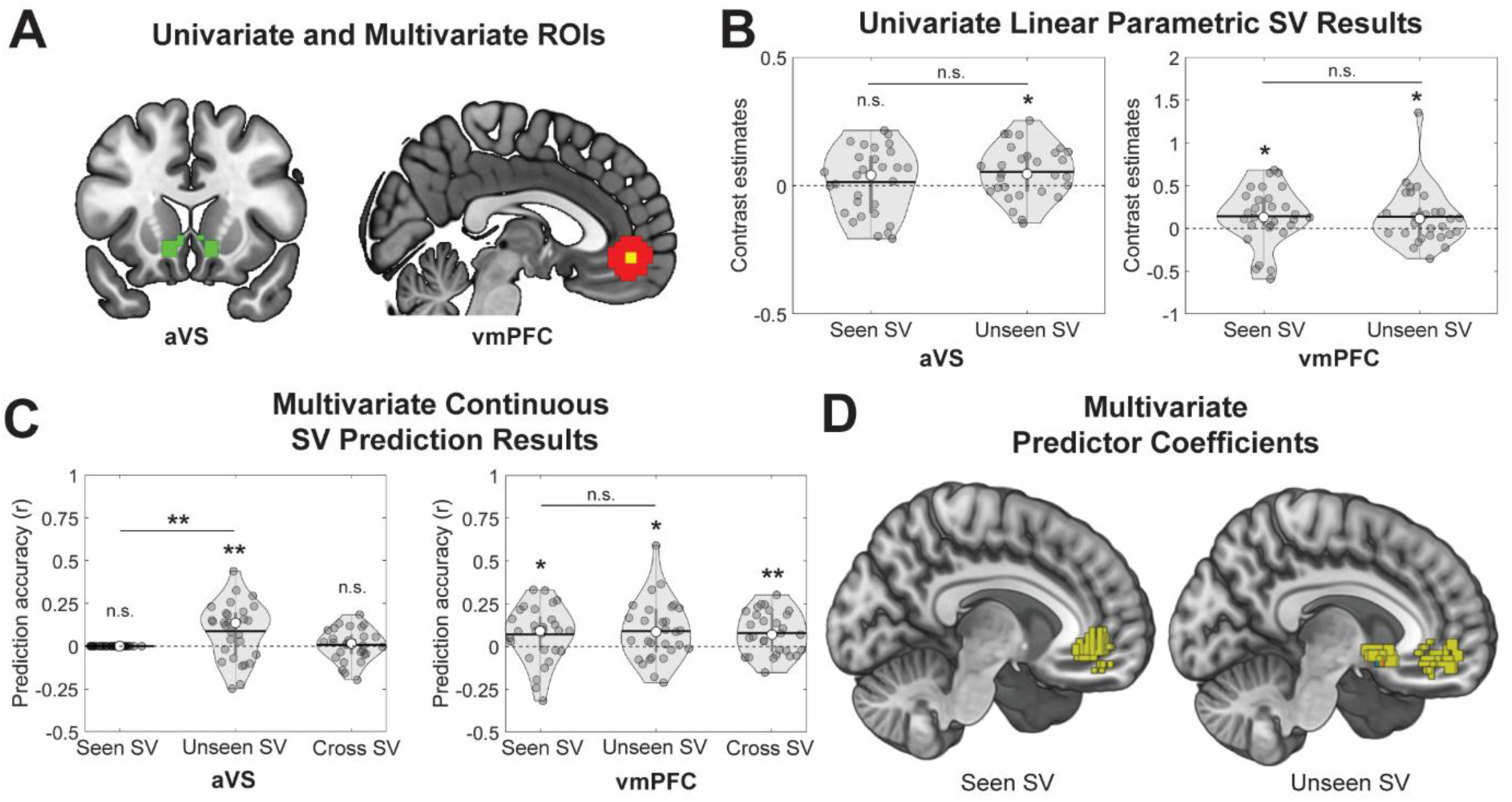
Subjective value (SV) processing. (**A**) Regions of interest (ROI) for univariate (yellow), multivariate (red), and both (green) analyses for the anterior ventral striatum (aVS) and ventromedial prefrontal cortex (vmPFC). (**B**) Univariate results for linear parametric SV contrasts for seen and unseen faces. (**C**) Multivariate continuous prediction results for SV from seen and unseen faces; and cross-decoding SV across seen and unseen faces. (**D**) Predictor coefficients for the subset of voxels that contributed to significant multivariate SV prediction results. Warm color indicates a positive, and cold color a negative, relationship with neural signal and higher SV. P-values were FDR-adjusted separately within each test set: 10 tests against chance and 4 paired tests. * p < .05, ** p ≤ .01, *** p ≤ .001.

##### Seen faces

In the vmPFC, SV ratings were positively related to mean neural signal (M = 0.14, SE = 0.06, t(28) = 2.26, p_FDR_ = .036, p_unc._ = .016, one-tailed), and multivariate regression with continuous SV ratings achieved significant out-of-sample prediction accuracy (Mdn_r_ = 0.091, p_FDR_ = .048, p_unc._ = .029, one-tailed permutation test), with positive predictor coefficients. In the aVS, however, no significant relationship was observed for mean BOLD signal (M = 0.01, SE = 0.02, t(28) = 0.57 p_FDR_ = .320, p_unc._ = .288, one-tailed) and prediction accuracy was non-significant (Mdn_r_ = 0, p_FDR_ = .822, p_unc._ = .822, one-tailed permutation test).

##### Unseen faces

In the vmPFC, SV ratings were again positively related to mean neural signal (M = 0.14, SE = 0.06, t(28) = 2.21, p_FDR_ = .036, p_unc._ = .018, one-tailed; Wilcoxon signed rank test to account for an outlier, Z = 2.00, p_FDR_ = .045, p_unc._ = .023, one-tailed), and multivariate regression produced significant prediction accuracy (Mdn_r_ = 0.086, p_FDR_ = .049, p_unc._ = .034, one-tailed permutation test) with positive predictor coefficients. In the aVS, SV ratings showed a significant positive relationship with mean neural signal (M = 0.05, SE = 0.02, t(28) = 2.80, p_FDR_ = .017, p_unc._ = .005, one-tailed), and multivariate regression significantly predicted SV (Mdn_r_ = 0.136, p_FDR_ = .010, p_unc._ = .002, one-tailed permutation test), with mostly positive predictor coefficients.

##### Cross-decoding

SV could be cross-decoded across seen and unseen faces in the vmPFC (Mdn_r_ = 0.069, p_FDR_ = .010, p_unc._ = .001, one-tailed permutation test), but not in the aVS (Mdn_r_ = 0.015, p_FDR_ = .151, p_unc._ = .121, one-tailed permutation test).

##### Seen and unseen face comparisons

In the vmPFC, the neural relationship with SV did not significantly differ in strength between seen and unseen faces with univariate (t(28) = 0.05, p_FDR_ = .999, p_unc._ = .962, two-tailed) or multivariate analysis (p_FDR_ > .999, p_unc._ > .999, two-tailed permutation test). In the aVS, the neural relationship with SV was significantly stronger for unseen than seen faces with multivariate (p_FDR_ = .012, p_unc._ = .003, two-tailed permutation test) but not with univariate analysis (t(28) = 1.22, p_FDR_ = .468, p_unc._ = .234, two-tailed). The results show that SV was processed at a similar strength in vmPFC when faces were seen and unseen, but at a greater strength in aVS when faces were unseen compared to seen, likely due to the lack of a significant neural SV-signal in aVS for seen faces.

Together, these findings suggest that SV from unseen faces was processed with a similar neural representation as SV from seen faces in vmPFC, but not in aVS, where only SV from unseen faces was detected.

#### Relationship between face perception in FFA and SV in aVS and vmPFC

We also examined whether face-signal strength in the FFA was related to SV-signal strength in the aVS and vmPFC (Figure 4). For unseen faces, Spearman rank correlations between FFA contrast estimates (unseen > absent) and SV-related parametric contrast estimates showed positive relationships with both aVS (r(27) = 0.43, p_FDR_ = .022, p_unc._ = .011, one-tailed) and vmPFC (r(27) = 0.57, p_FDR_ = .004, p_unc._ = .001, one-tailed). For seen faces, however, no significant relationships were found with aVS (r(27) = 0.19, p_FDR_ = .213, p_unc._ = .160, one-tailed) or vmPFC (r(27) = −0.08, p_FDR_ = .332, p_unc._ = .332, one-tailed). These findings suggest that stronger unseen face processing in the FFA is associated with stronger SV processing in both aVS and vmPFC.

**Figure 4.**
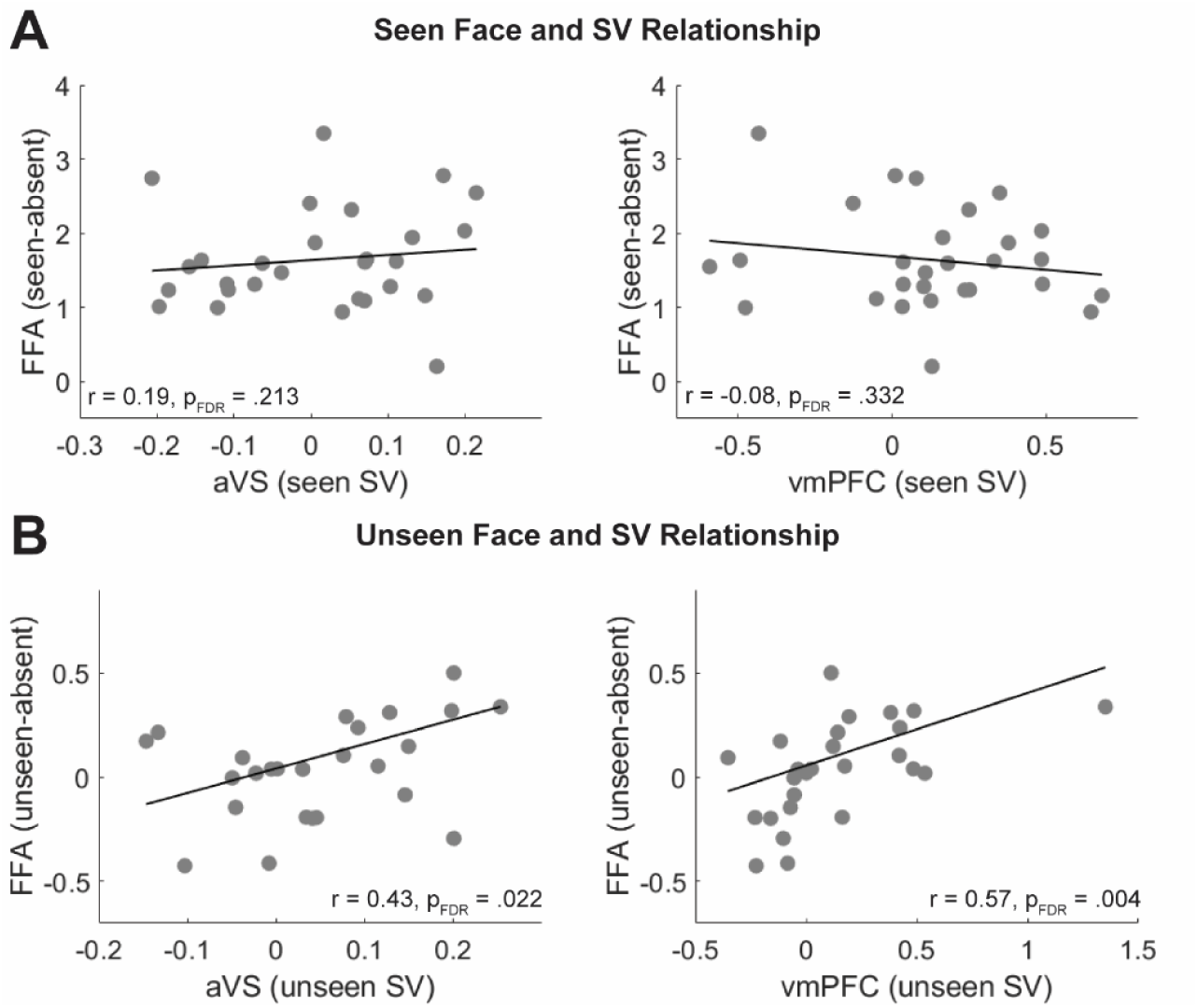
Relationship between face and SV signal. Univariate TFCE adjusted and FDR corrected (p < 0.05, one-tailed) contrasts showing linear parametric results for subjective value (SV) for seen faces, seen > absent face presentation, seen > absent delay, seen > absent response. None of the whole-brain contrasts with unseen faces survived corrections.

#### Relationship between neural SV signals and behavior

There was no significant relationships between SV processing from seen faces in aVS (r(27) = 0.20, p_FDR_ = .766, p_unc._ = .304, two-tailed) or vmPFC (r(27) = −0.06, p_FDR_ = . 766, p_unc._ = .766, two-tailed) and reaction time. Similarly, we did not find any significant relationship between SV processing from unseen faces in aVS (r(27) = 0.09, p_FDR_ = . 766, p_unc._ = .660, two-tailed) or vmPFC (r(27) = 0.07, p_FDR_ = . 766, p_unc._ = .701, two-tailed) and reaction time.

#### Whole-brain results

TFCE-adjusted and FDR corrected whole-brain contrasts were used to explore whether there was neural signal in any brain areas outside of our ROIs that was associated with seen or unseen face perception and seen or unseen SV processing.

##### Seen faces

Seen > absent faces revealed significant neural signal in the bilateral vmPFC and right aVS, together with many other brain areas, including bilateral dorsal posterior cingulate cortex (dPPC), precuneus, and large bilateral areas of the temporal cortex (including FFA), parietal cortex, frontal cortex, cerebellum, and subcortical areas such as putamen, amygdala, and hippocampus (SI-Figure 3). The linear parametric SV modulation of seen faces revealed several clusters where neural signal had a positive relationship to SV, including bilaterally in the anterior cingulate cortex and extending through the dorsomedial prefrontal cortex (dmPFC), the cerebellum, the middle frontal gyrus, and precentral gyrus (SI-Figure 3).

##### Unseen faces

Results for unseen > absent and parametric SV modulations of unseen faces did not survive FDR corrections. Uncorrected results for unseen faces are provided in the Supplementary Information (SI-Figure 4-5).

##### Seen and unseen face comparisons

Seen > unseen faces did not reveal any difference in the vmPFC. However, seen faces were associated with a greater neural signal in the right aVS and several other brain areas, including bilateral dPPC, precuneus, and large bilateral areas of the temporal cortex (including FFA), parietal cortex, frontal cortex, cerebellum, and subcortical areas such as putamen, amygdala, and hippocampus (SI-Figure 3). Instead, unseen faces were associated with a greater neural signal in areas such as the bilateral occipital cortex, supramarginal gyrus, inferior frontal gyrus and precentral gyrus, ventral posterior cingulate cortex (vPPC), precuneus, insula, and subcortical areas such as caudate nucleus (extending slightly into left aVS) and thalamus (SI-Figure 3). When comparing the parametric SV contrasts for seen and unseen faces, no differences survived FDR corrections (for uncorrected results, see SI-Figure 4-5).

##### Relationship between neural SV signal and behavior

Neither whole brain correlations between seen reaction times and neural SV signals from seen faces, or correlations between unseen reaction times and neural SV signals from unseen faces, survived whole-brain FDR corrections for multiple comparisons (for uncorrected results, see SI-Figure 6).

## Discussion

We tested whether the subjective value (SV) of faces, captured by idiosyncratic ratings of overall facial attractiveness, can be processed automatically in the aVS and vmPFC without conscious awareness. We found that (i) subliminal faces elicited a similar neural SV representation as that of conscious faces in vmPFC, but not in aVS, where only unseen faces elicited a detectable SV response; (ii) subliminal faces were processed in FFA with a similar but weaker neural representation as conscious faces; and (iii) stronger subliminal face processing in FFA was associated with stronger subliminal SV processing in aVS and vmPFC. These results suggest that SV can be computed automatically in a bottom-up fashion in core valuation circuits even when the stimulus is outside conscious awareness.

Our findings provide strong evidence that SV can be computed automatically in core valuation circuits even when stimuli are entirely outside awareness, extending previous work on automatic SV computation by addressing limitations in earlier studies. Previous studies have shown that aVS and vmPFC can encode SV during passive viewing or unrelated tasks (Kim et al., 2007; Lebreton et al., 2009; Levy et al., 2011; Smith et al., 2014; Tusche et al., 2010), but it is difficult to rule out spontaneous explicit evaluations when stimuli are consciously presented. Subliminal presentations remove this confound, but previous attempts to probe SV under such conditions were limited in several ways (Bijleveld et al., 2014; Fernandes et al., 2025; Gillath & Canterberry, 2012; Ito et al., 2015; Oei et al., 2012; Pessiglione et al., 2007, 2008): (i) trial-wise measures of awareness were not used, allowing for the possibility of conscious trials being included and influencing the subliminal results; (ii) SV was not measured with idiosyncratic participant-specific ratings; (iii) subliminal information was often relevant to the task so top-down influences could not be ruled out; and/or (iv) SV effects were not found in the brain’s valuation areas (aVS and vmPFC). We addressed all of these issues by combining CFS with trial-wise objective and subjective awareness measures in an MRI scanner, and directly linked neural signal in core valuation areas to participant-specific continuous post-fMRI SV ratings. Our results therefore provide the strongest evidence to date that core valuation circuits, including the vmPFC, can automatically compute SV without conscious awareness, and we demonstrate this for the first time using facial attractiveness.

The finding that subliminal SV processing extended from aVS to vmPFC is consistent with our earlier work showing that subliminal probabilities can be integrated with consciously perceived rewards into SV within these same regions (Fernandes et al., 2025). Together, these two studies, which used different stimuli and tasks, make a strong case that SV computations can operate outside conscious awareness. The evidence supports the idea that facial attractiveness engages the same domain-general “common currency” circuitry as other value domains. From an evolutionary standpoint, facial attractiveness is likely a form of SV that reflects the desirability of a potential mate that influences mate selection (Buss, 1989; Buss & Schmitt, 2019). This is further supported by univariate studies and meta-analyses showing overlapping aVS and vmPFC signals for faces and other rewards (Bartra et al., 2013; Chuan-Peng et al., 2020; Kim et al., 2007; Lebreton et al., 2009), and by findings that neural representations of attractiveness generalize across stimulus categories (Pegors et al., 2015). Our results therefore challenge the prevailing assumption that subliminal SV processing is confined to rudimentary subcortical mechanisms (Bijleveld et al., 2012), and add to previous reports of subliminal value processing in subcortical regions (Fernandes et al., 2025; Oei et al., 2012; Pessiglione et al., 2007, 2008). However, in this and previous studies, the subliminal stimuli represented a one-dimensional SV (e.g., facial attractiveness, probability, or monetary reward). It remains an open question whether subliminal SV also can be automatically computed from more complex stimuli where the overall SV is a composite of many different value-dimensions or whether several value-dimensions can be integrated into SV without conscious awareness.

Our results stand in contrast to the only other fMRI study on subliminal facial attractiveness, which only found aVS and vmPFC effects for consciously seen faces (Ito et al., 2015). Methodological differences likely explain this discrepancy. For example, their brief backward-masked presentations (34 ms) may have produced weaker subliminal effects than our three-second CFS presentations, and their ROI definition may have missed the vmPFC subregion most reliably associated with SV and facial attractiveness (Bartra et al., 2013; Chuan-Peng et al., 2020). Instead, subliminal attractiveness effects have been reported outside valuation circuits, such as marginal effects in dmPFC, ventrolateral prefrontal cortex/anterior insula (Ito et al., 2015), and posterior visual/parietal areas measured with EEG (Hou et al., 2023; Shang et al., 2025). These regions are more commonly associated with salience, the ability of a stimulus to capture attention or increase general arousal, rather than SV per se.

Salience has a U-shaped relationship with SV but will correlate with SV when negative values are absent, and is typically associated with neural signal in areas such as the anterior insula, caudate nucleus, dmPFC, dorsolateral prefrontal cortex, and inferior posterior parietal cortex (Bartra et al., 2013; Kahnt et al., 2014; Kahnt & Tobler, 2017; Litt et al., 2011; Zink et al., 2004). While our design cannot dissociate SV from salience because our stimuli lacked negatively rated faces, prior work shows that when these factors are dissociated, signals in aVS and vmPFC reflects SV but not salience (Bartra et al., 2013; Fernandes et al., 2025; Kahnt et al., 2014; Litt et al., 2011). Our whole-brain results showing an association between neural signal from seen faces and SV ratings in the dmPFC may therefore reflect a salience rather than SV signal.

We also replicate earlier findings of subliminal face processing in the FFA (Fang & He, 2005; Jiang & He, 2006; Moutoussis & Zeki, 2002). Notably, we found that stronger subliminal face responses in FFA correlated with stronger SV responses in aVS and vmPFC. The FFA is specialized for processing invariant facial structures important for identification (George et al., 1999; Hoffman & Haxby, 2000; Kanwisher et al., 1997) and likely provides the structural information on which attractiveness judgments rely, such as symmetry, averageness, and sexual dimorphism (Rhodes, 2006). This interpretation is consistent with evidence from effective connectivity from FFA to vmPFC (Fairhall & Ishai, 2007) and with clinical findings that prosopagnosia patients with FFA lesions have impaired attractiveness judgments (Iaria et al., 2008). The absence of this correlation for seen faces may indicate that the relationship is more apparent when processing is strongly suppressed, with suppression strength varying across individuals.

We did not find SV signals in aVS for conscious faces, despite robust effects for subliminal faces. One possibility is that, for seen faces, task-related neural signals may have masked SV signals in aVS, and perhaps to a lesser extent in vmPFC. Consciously seen faces likely engaged the task differently than subliminal faces, which may have been processed more passively. Additionally, because faces were absent on most trials, their sudden appearance could have triggered intrinsic reward predictions or motivational signals related to anticipating successful task performance. Although such effects have not been examined prior to decision-making, there is evidence for intrinsic reward prediction errors in aVS during successful memory performance without task feedback or incentives (Satterthwaite et al., 2012; Wolf et al., 2011). Such signals might have interfered with SV signals and contributed to weaker effects for seen faces in our experiment.

Although subliminal faces did not directly influence task performance, participants responded more slowly to subliminal than absent trials, suggesting a lingering memory trace. This is consistent with previous results from similar memory paradigms (Bergström & Eriksson, 2014, 2015, 2018; Pedale et al., 2023). It does not seem to be the case that SV processing in aVS or vmPFC had an influence on reaction time for seen or unseen faces, which could be because the face-identification task was not directly related to SV. However, it is worth noting that the uncorrected whole-brain results revealed a trend that SV processing in vPCC from unseen and seen faces correlated positively with reaction time, similar to findings from Grueschow et al., (2015).

A continuing debate in subliminal cognition concerns how to best measure the absence of conscious awareness. Subjective measures best capture the subjective experience itself but may overestimate unawareness by misclassifying weak conscious signals as unseen. Objective measures classify any above-chance performance as conscious awareness, which may underestimate unawareness by misclassifying unconscious signals as conscious. Importantly, however, the two approaches often converge on similar findings across the literature (Dehaene & Changeux, 2011). In our study, we used both subjective (PAS) and objective (face identity) trial-wise measures. Participants were trained to report PAS ratings using two buttons (1 = no visual experience, 2 = vague-to-clear visual experience; Fernandes et al., 2025), which was sufficient for our purpose of distinguishing between “no experience” and “vague visual experience.” Suppressed faces and absent trials were both consistently rated as “no experience,” and objective performance was at chance for unseen faces. The fact that we observed reaction time, neural face, and neural SV effects from unseen faces when both subjective and objective trial-wise measures indicated unawareness, strongly supports that these effects were truly based on subliminal processing.

In conclusion, we show that SV (operationalized as facial attractiveness) is automatically processed in aVS and vmPFC without the need for conscious awareness, demonstrating that stimulus value processing can occur independently of conscious awareness. These results suggest an avenue for automatic subliminal influences on important value-based decisions (e.g., mate selection). Future research will have to investigate whether conscious awareness is necessary for later stages in the decision process (e.g., action value integration from stimulus value and action cost).

## Author Contributions

FB: Conceptualization, funding acquisition, supervision, project administration, methodology, investigation, formal analysis, visualization, validation, data curation, writing – original draft, writing – review & editing. PF: Investigation, formal analysis, writing – review & editing. JWK: Methodology and writing – review & editing. AT: Methodology and writing – review & editing. JA: Methodology and writing – review & editing. CR: Methodology and writing – review & editing.

## Declaration of Competing Interest

The authors declare no competing interests.

## Code availability statement

The Matlab code for the SHINE Toolbox can be found at Github (https://github.com/RodDalBen/SHINE_color). The custom Matlab code used to create the CFS stimuli can be found at Martin Hebart’s webpage (http://martin-hebart.de/webpages/code/stimuli.html). The Thresholded Partial Least Squares (T-PLS) toolbox used for the multivariate decoding analyses can be found at Github (https://github.com/sangillee/TPLSm).

## Funding/Acknowledgements

This work was supported by The BIAL Foundation (A-27942 to FB). FB was supported by Fundação para a Ciência e Tecnologia (CEECIND/03661/2017). JA was supported by a European Research Counsil (ERC) under the European Union’s Horizon 2020 research and innovation program Starting Grant number 802553 “Content Map”, and by European Research Executive Agency Widening program under the European Union’s Horizon Europe Grant 101087584 “Cogbooster”.

## Notes

### Competing Interest Statement

The authors have declared no competing interest.

