## Supplemental Information for "Subliminal beauty engages the brain’s valuation circuits"

### SI-Methods

#### fMRIprep preprocessing biolerplate

Results included in this manuscript come from preprocessing performed using fMRIPrep 24.0.1 (Esteban et al. (2019); Esteban et al. (2018); RRID:SCR_016216), which is based on Nipype 1.8.6 (K. Gorgolewski et al. (2011); K. J. Gorgolewski et al. (2018); RRID:SCR_002502).

*Anatomical data preprocessing*

A total of 1 T1-weighted (T1w) images were found within the input BIDS dataset. The T1w image was corrected for intensity non-uniformity (INU) with N4BiasFieldCorrection (Tustison et al. 2010), distributed with ANTs 2.5.1 (Avants et al. 2008, RRID:SCR_004757), and used as T1w-reference throughout the workflow. The T1w-reference was then skull-stripped with a Nipype implementation of the antsBrainExtraction.sh workflow (from ANTs), using OASIS30ANTs as target template. Brain tissue segmentation of cerebrospinal fluid (CSF), white-matter (WM) and gray-matter (GM) was performed on the brain-extracted T1w using fast (FSL (version unknown), RRID:SCR_002823, Zhang, Brady, and Smith 2001). Brain surfaces were reconstructed using recon-all (FreeSurfer 7.3.2, RRID:SCR_001847, Dale, Fischl, and Sereno 1999), and the brain mask estimated previously was refined with a custom variation of the method to reconcile ANTs-derived and FreeSurfer-derived segmentations of the cortical gray-matter of Mindboggle (RRID:SCR_002438, Klein et al. 2017). Volume-based spatial normalization to one standard space (MNI152NLin2009cAsym) was performed through nonlinear registration with antsRegistration (ANTs 2.5.1), using brain-extracted versions of both T1w reference and the T1w template. The following template was were selected for spatial normalization and accessed with TemplateFlow (24.2.0, Ciric et al. 2022): ICBM 152 Nonlinear Asymmetrical template version 2009c [Fonov et al. (2009), RRID:SCR_008796; TemplateFlow ID: MNI152NLin2009cAsym].

*Functional data preprocessing*

For each of the 3 BOLD runs found per subject (across all tasks and sessions), the following preprocessing was performed. First, a reference volume was generated, using a custom methodology of fMRIPrep, for use in head motion correction. Head-motion parameters with respect to the BOLD reference (transformation matrices, and six corresponding rotation and translation parameters) are estimated before any spatiotemporal filtering using mcflirt (FSL, Jenkinson et al. 2002). The BOLD reference was then co-registered to the T1w reference using bbregister (FreeSurfer) which implements boundary-based registration (Greve and Fischl 2009). Co-registration was configured with six degrees of freedom. Several confounding time-series were calculated based on the preprocessed BOLD: framewise displacement (FD), DVARS and three region-wise global signals. FD was computed using two formulations following Power (absolute sum of relative motions, Power et al. (2014)) and Jenkinson (relative root mean square displacement between affines, Jenkinson et al. (2002)). FD and

DVARS are calculated for each functional run, both using their implementations in Nipype (following the definitions by Power et al. 2014). The three global signals are extracted within the CSF, the WM, and the whole-brain masks. Additionally, a set of physiological regressors were extracted to allow for component-based noise correction (CompCor, Behzadi et al. 2007). Principal components are estimated after high-pass filtering the preprocessed BOLD time-series (using a discrete cosine filter with 128s cut-off) for the two CompCor variants: temporal (tCompCor) and anatomical (aCompCor). tCompCor components are then calculated from the top 2% variable voxels within the brain mask. For aCompCor, three probabilistic masks (CSF, WM and combined CSF+WM) are generated in anatomical space. The implementation differs from that of Behzadi et al. in that instead of eroding the masks by 2 pixels on BOLD space, a mask of pixels that likely contain a volume fraction of GM is subtracted from the aCompCor masks. This mask is obtained by dilating a GM mask extracted from the FreeSurfer's aseg segmentation, and it ensures components are not extracted from voxels containing a minimal fraction of GM. Finally, these masks are resampled into BOLD space and binarized by thresholding at 0.99 (as in the original implementation). Components are also calculated separately within the WM and CSF masks. For each CompCor decomposition, the k components with the largest singular values are retained, such that the retained components' time series are sufficient to explain 50 percent of variance across the nuisance mask (CSF, WM, combined, or temporal). The remaining components are dropped from consideration. The head-motion estimates calculated in the correction step were also placed within the corresponding confounds file. The confound time series derived from head motion estimates and global signals were expanded with the inclusion of temporal derivatives and quadratic terms for each (Satterthwaite et al. 2013). Frames that exceeded a threshold of 0.5 mm FD or 1.5 standardized DVARS were annotated as motion outliers. Additional nuisance timeseries are calculated by means of principal components analysis of the signal found within a thin band (crown) of voxels around the edge of the brain, as proposed by (Patriat, Reynolds, and Birn 2017). All resamplings can be performed with a single interpolation step by composing all the pertinent transformations (i.e. head-motion transform matrices, susceptibility distortion correction when available, and co-registrations to anatomical and output spaces). Gridded (volumetric) resamplings were performed using nitransforms, configured with cubic B-spline interpolation.

Many internal operations of fMRIPrep use Nilearn 0.10.4 (Abraham et al. 2014, RRID:SCR_001362), mostly within the functional processing workflow. For more details of the pipeline, see the section corresponding to workflows in fMRIPrep's documentation.

*Copyright Waiver*

The above boilerplate text was automatically generated by fMRIPrep with the express intention that users should copy and paste this text into their manuscripts unchanged. It is released under the CC0 license.

**SI-Methods Figure**

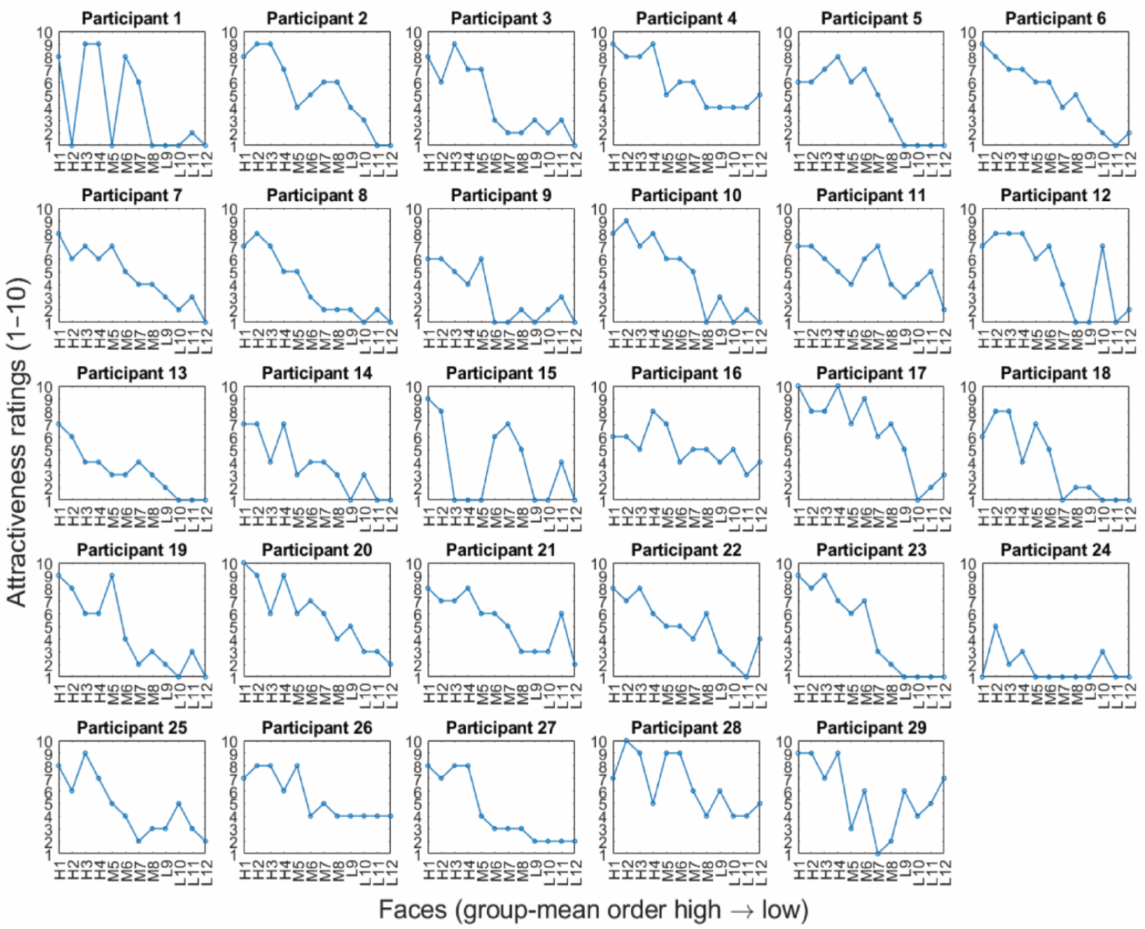


**SI-Figure 1. Individual attractiveness ratings.** Plots show the individual attractiveness ratings (1-10) with face images ordered from highest to lowest group mean ratings, and shows each individual's idiosyncratic attractiveness ratings. Face image ID denotes the number from highest to lowest group mean attractiveness ratings and what predetermined attractiveness group they belonged to (High, Medium, or Low).

### SI-Results

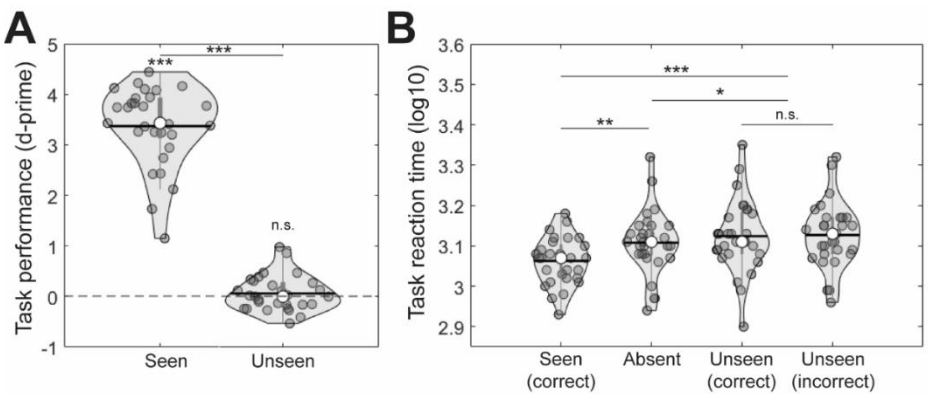


**SI-Figure 2. Behavioral results.** (A) Face-identification task performance (d-prime) for seen and unseen faces (uncorrected p-values). (B) Face identification reaction time (log10-transformed) for seen correct, absent, unseen correct, and unseen incorrect trials (FDR adjusted p-values for 6 pairwise tests). * p < .05, ** p ≤ .01, *** p ≤ .001.

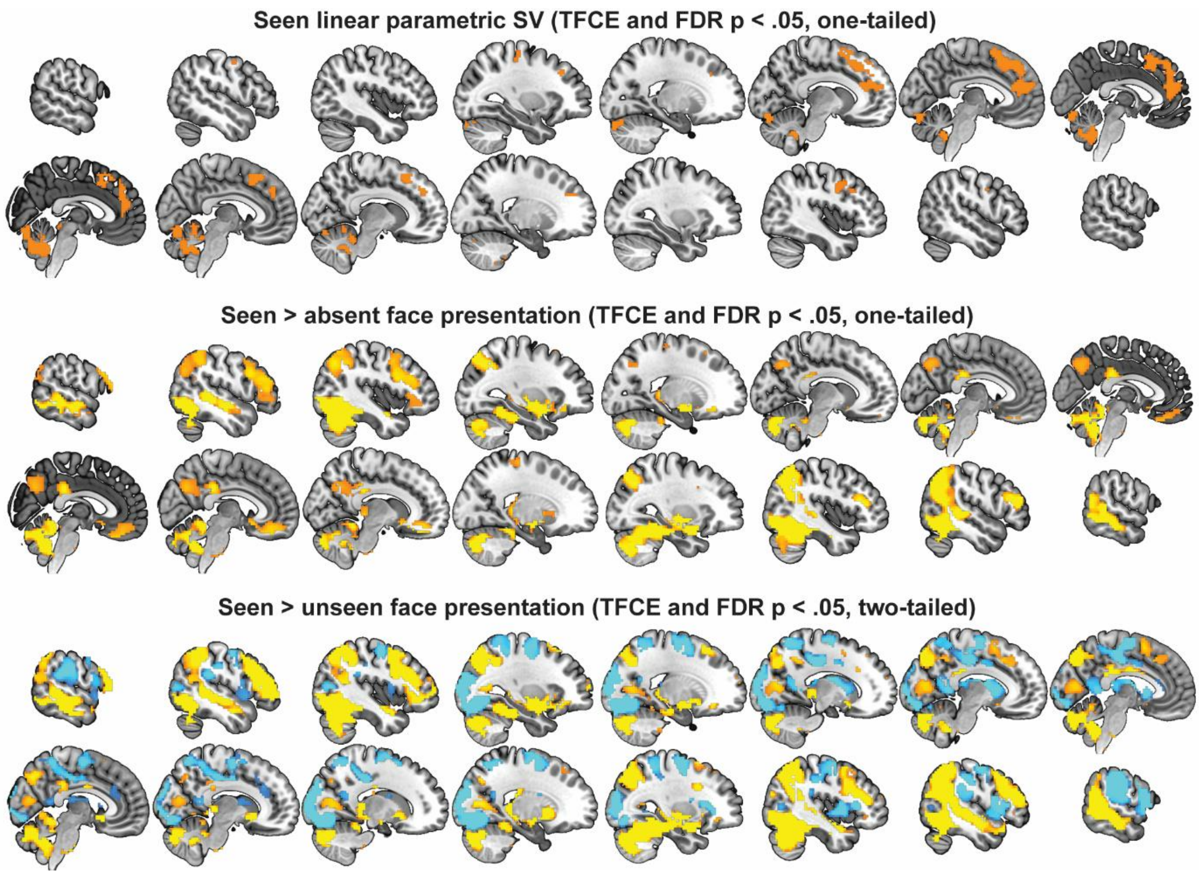

**SI-Figure 3. TFCE and FDR corrected whole-brain results.** Univariate threshold free cluster enhancement (TFCE) adjusted and FDR corrected ($p < 0.05$, one-tailed) contrasts: (i) Linear parametric modulation results for subjective value (SV) of seen faces. (ii) Seen > absent face presentations. (iii) Seen > unseen face presentations. None of the whole-brain contrasts with unseen faces survived corrections. Warm colors are positive and cold colors are negative associations.

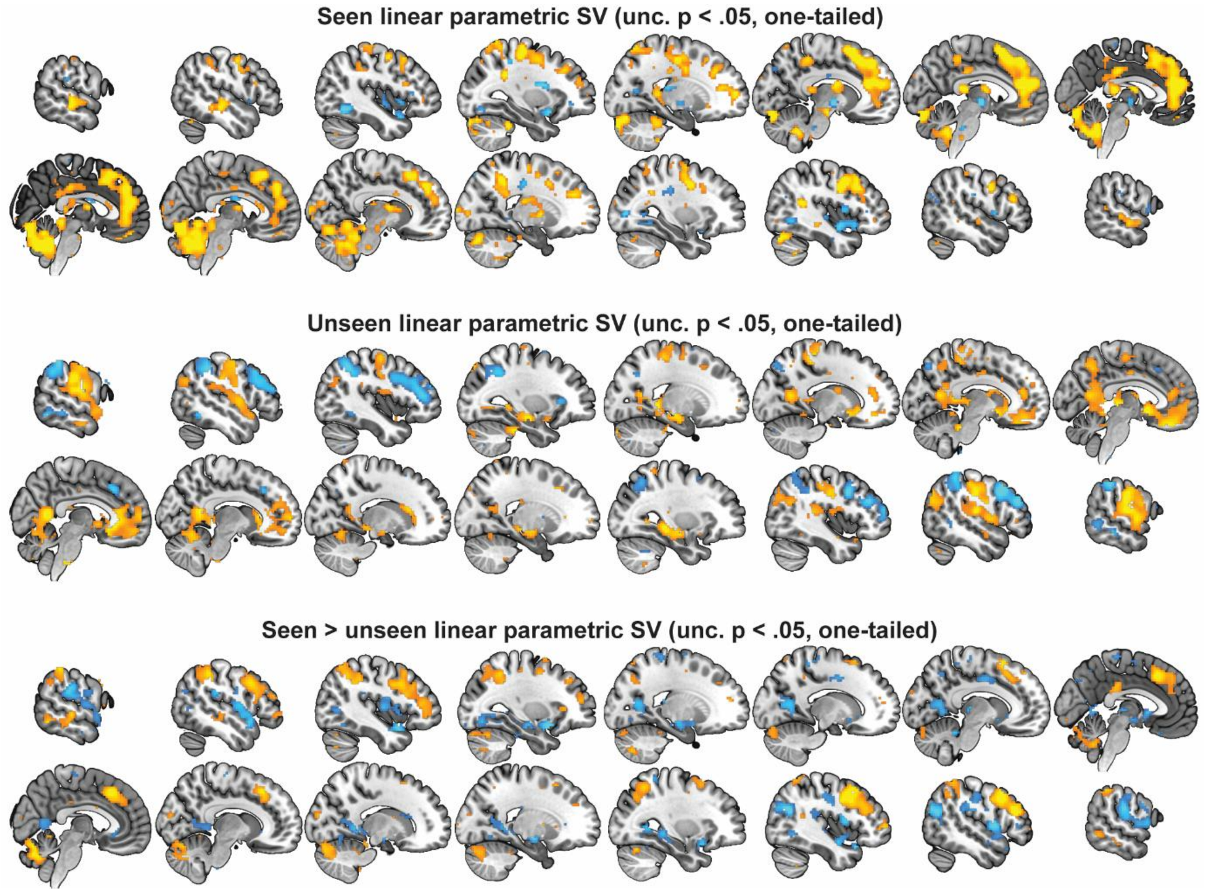


**SI-Figure 4. Uncorrected whole-brain results for parametric SV modulation.** Uncorrected univariate contrasts ($p < 0.05$, one-tailed): (i) Linear parametric modulation results for subjective value (SV) of seen faces. (ii) Linear parametric modulation results for SV of unseen faces. (iii) Seen SV > unseen SV. Warm colors are positive and cold colors are negative associations.

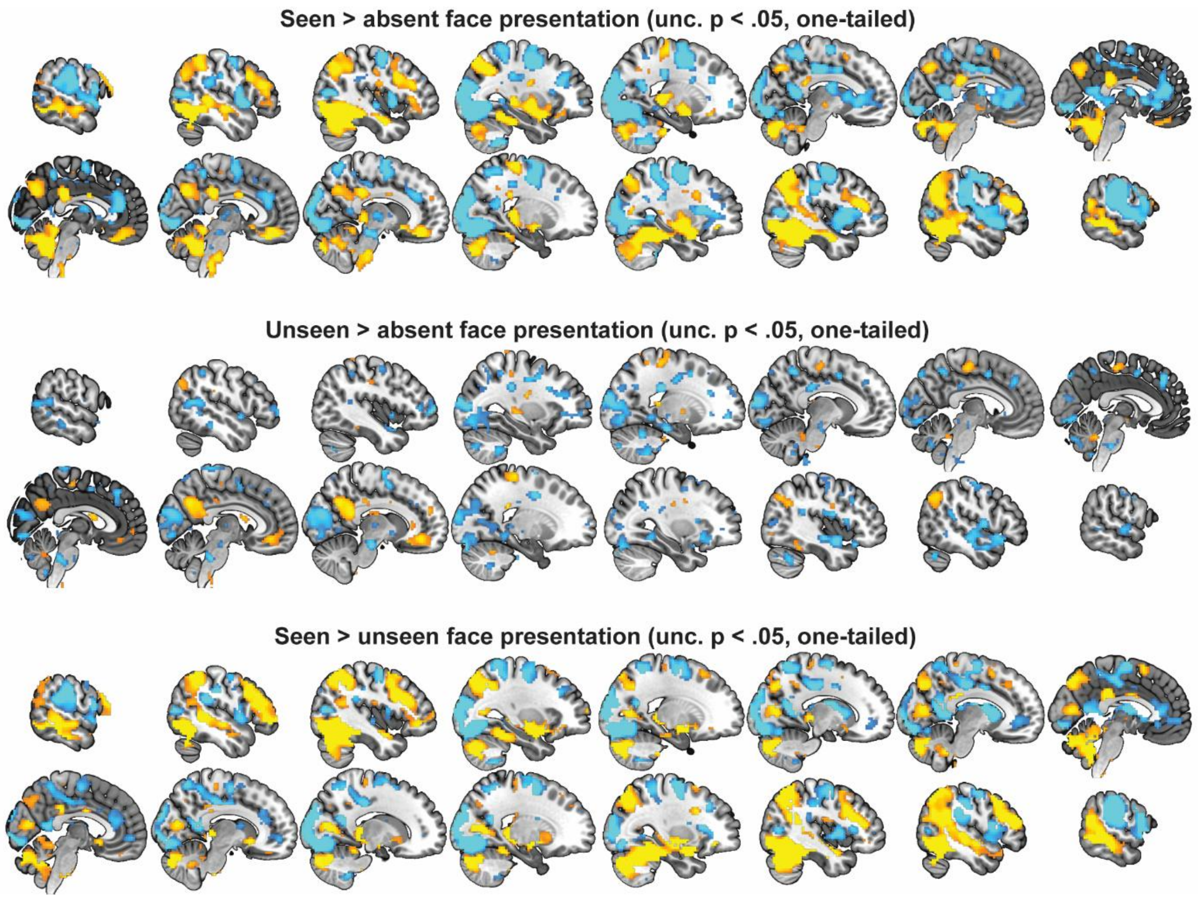


**SI-Figure 5. Uncorrected whole-brain results for seen, unseen, and absent faces.** Uncorrected univariate contrasts ($p < 0.05$, one-tailed): (i) Seen > absent face presentations. (ii) Unseen > absent face presentations. (iii) Seen > unseen face presentations. Warm colors are positive and cold colors are negative associations.

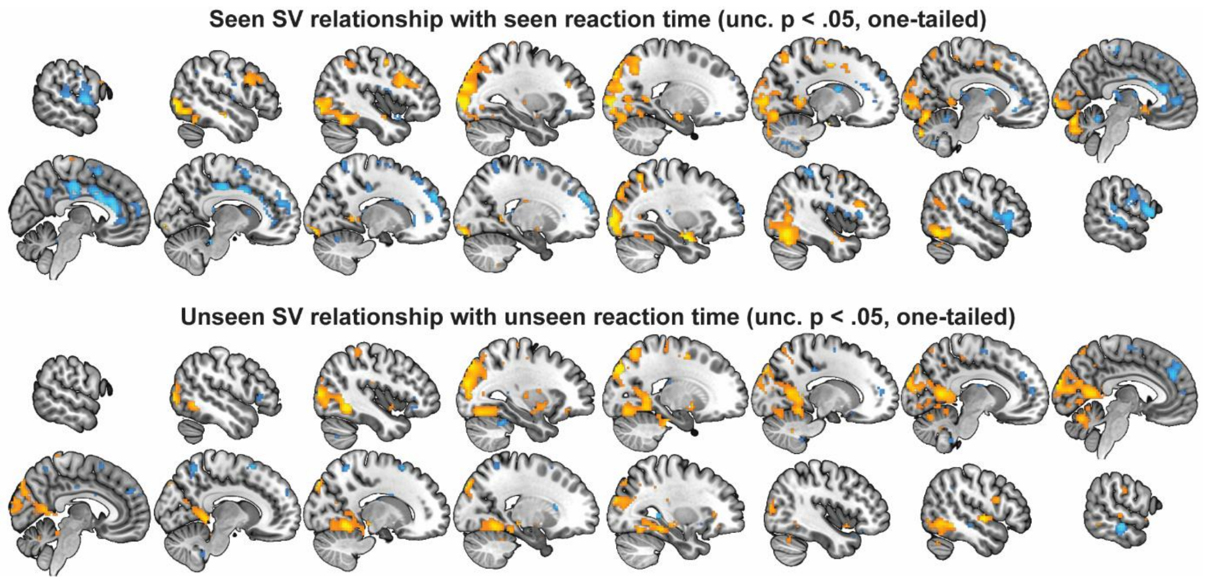


**SI-Figure 6. Uncorrected whole-brain results for relationship between parametric SV and reaction times.** Uncorrected univariate contrasts ($p < 0.05$, one-tailed): (i) Correlations between parametric SV contrast estimates from seen faces and seen reaction time. (ii) Correlations between parametric SV contrast estimates from seen faces and seen reaction time. Warm colors are positive and cold colors are negative associations.
